# RPEGENE-Net: A Multi-Resolution Deep Learning Framework for Predicting Gene Expression from Microscopy Images of Retinal Pigment Epithelium (RPE) Cells

**DOI:** 10.1101/2025.08.27.672622

**Authors:** M. Hossein Nowroozzadeh, Neda Taghinezhad, Tahereh Mahmoudi, Fatemeh Sanie-Jahromi

**Author notes:** **Corresponding author:** Tahereh Mahmoudi, Fatemeh Sanie-Jahromi (same correspondence) Postal address: Zand Boulevard, Poostchi Street, Poostchi Ophthalmology Research Center, Shiraz, Iran., Tahereh Mahmoudi. **Authors email:** M. Hossein Nowroozzadeh, Neda Taghinezhad.

## Abstract

**Purpose:** To develop a deep learning framework, RPEGENE-Net, capable of predicting gene expression profiles of retinal pigment epithelium (RPE) cells using live-cell microscopy images.

**Methods:** A dataset of live-cell images of RPE cells, treated with various drug regimens and captured at magnifications of 40x, 100x, 200x, and 400x, was used. Gene expression of six key genes involved in epithelial-mesenchymal transition or EMT (including α-SMA, ZEB1, TGF-β, CD90, β-catenin, Snail) and treatment classes (Aflibercept, Bevacizumab, Dexamethasone, Aflibercept + Dexamethasone, and untreated control) were analyzed. After preprocessing the image data and gene expression values, we trained and evaluated twelve state-of-the-art deep learning architectures, including three variants of DenseNet, five variants of ResNet, EfficientNet_b5, Inception_v3, RegNet_y_400mf, and a vision transformer model (Swin_b). A two-stage pipeline was implemented, combining autoencoder-based pretraining to extract meaningful features with fine-tuning specifically optimized for gene expression regression and treatment type classification tasks. Features extracted from the second stage across four magnifications were concatenated to generate the final prediction, leveraging multi-scale morphological information for improved accuracy.

**Results:** DenseNet121 demonstrated superior performance, achieving the highest Pearson correlation coefficients for four genes: α-SMA (0.79), ZEB1(0.84), TGF-β (0.83), and Snail (0.86). ResNet34 outperformed other models for CD90 (0.87) and β-catenin (0.85) predictions. The average mean absolute error (MAE) and average root mean square error (RMSE) on test dataset were 0.0244 and 0.1228, respectively. The R^2^ scores ranged from 0.50 (α-SMA) to 0.74 (TGF-β), indicating strong alignment between predicted and actual gene expression values. A multi-level approach, combining data from 40x, 100x, 200x, and 400x magnifications yielded higher R^2^ scores for almost all genes compared to single-magnification models. For the classification task, DenseNet121 achieved F1 score, precision, recall, and accuracy of 0.98, with a specificity of 0.99.

**Conclusions:** RPEGENE-Net provides a simple, cost-effective method to predict gene expression from live-cell images, with potential applications in experimental studies, RPE transplantation quality control, and broader cell-based research. Multi-magnification imaging enhances model performance, supporting its utility as a scalable tool for diverse gene expression studies.

## 1. Introduction

Retinal pigment epithelium (RPE) cells play a pivotal role in maintaining retinal homeostasis and are central to the pathogenesis of retinal degenerative diseases such as age-related macular degeneration (AMD) and proliferative vitreoretinopathy (PVR). In vitro RPE cell models have become indispensable tools for studying disease mechanisms, including oxidative stress, inflammation, and cell death, as well as for evaluating the efficacy, toxicity, and safety of therapeutic interventions [1, 2]. These models are also critical for advancing RPE transplantation research, enabling the refinement of cell culture techniques and the assessment of transplanted cell viability and functionality [3]. Commonly used RPE models include spontaneously formed cell lines (e.g., ARPE-19)[4], immortalized lines (e.g., hTERT-RPE1, RPE-J, D407)[5], primary human or animal RPE cells, and embryonic or induced pluripotent stem cell (iPSC)-derived RPE cells [6].

Gene expression analysis is a cornerstone of cellular research, providing critical insights into protein expression and cellular behavior. Techniques such as quantitative real-time PCR (qPCR), microarrays, and RNA sequencing have been widely used to study RPE cells, offering valuable data for understanding disease mechanisms and identifying therapeutic targets [7, 8]. However, these methods are often limited by high costs, the need for specialized equipment, and reliance on skilled operators [9, 10]. Furthermore, preparatory procedures such as cell lysis, fixation, and staining—essential for techniques like PCR and flow cytometry—render the cells unsuitable for further clinical use, limiting their application in final quality control during clinical phases [11]. These challenges highlight the need for non-invasive, cost-effective alternatives for gene expression analysis.

Emerging evidence suggests a potential link between gene expression and cell morphology, offering a novel approach to predict gene expression patterns without compromising cell viability [12, 13]. Cell morphology can be directly influenced by cytoskeletal genes and cell adhesion molecules or indirectly modulated by metabolic genes, signaling pathways (e.g., Wnt, Notch), and epigenetic regulation [14]. Although these morphological changes are often subtle, they can provide valuable cues for predicting gene expression, enabling the analysis of cellular responses to therapies or manipulations in a non-destructive manner. Recent studies have demonstrated the feasibility of this approach in various cell types, paving the way for its application in RPE research [15–17].

In this study, we introduce RPEGENE-Net, a multi-resolution deep learning framework designed to predict gene expression in RPE cells based on microscopy images. The model focuses on six key genes involved in epithelial-mesenchymal transition (EMT)—α-SMA, ZEB1, TGF-β, CD90, β-catenin, and Snail—and five treatment classes: aflibercept, bevacizumab, dexamethasone, a combination of aflibercept and dexamethasone, and a control group with no drug treatment. EMT is a critical process in RPE pathology, contributing to fibrosis and vision loss in diseases such as AMD and PVR [18]. The selected genes are well-established markers of EMT, regulating cytoskeletal remodeling, cell migration, and extracellular matrix deposition [19]. The chosen drugs include anti-VEGF agents (aflibercept, bevacizumab) and an anti-inflammatory corticosteroid (dexamethasone), which target key pathways implicated in RPE dysfunction [20].

By leveraging advanced deep learning techniques, RPEGENE-Net aims to provide a non-invasive, high-throughput method for predicting gene expression in RPE cells, offering a powerful tool for both basic research and clinical applications.

## 2. Materials and Methods

This study was approved by the Ethics Committee of Shiraz University of Medical Sciences (IR.SUMS.REC.1401.458). All methods were carried out in accordance with relevant guidelines and regulations. Figure 1 illustrates the core concept of the present study: extracting gene expression profiles from cell morphology data. This approach encompasses both static and dynamic methodologies (with only the static approach explored in this study), each addressing distinct aspects of cell morphology. We propose that integrating models across these features could enhance the accuracy of the final composite model.

**Figure 1.**
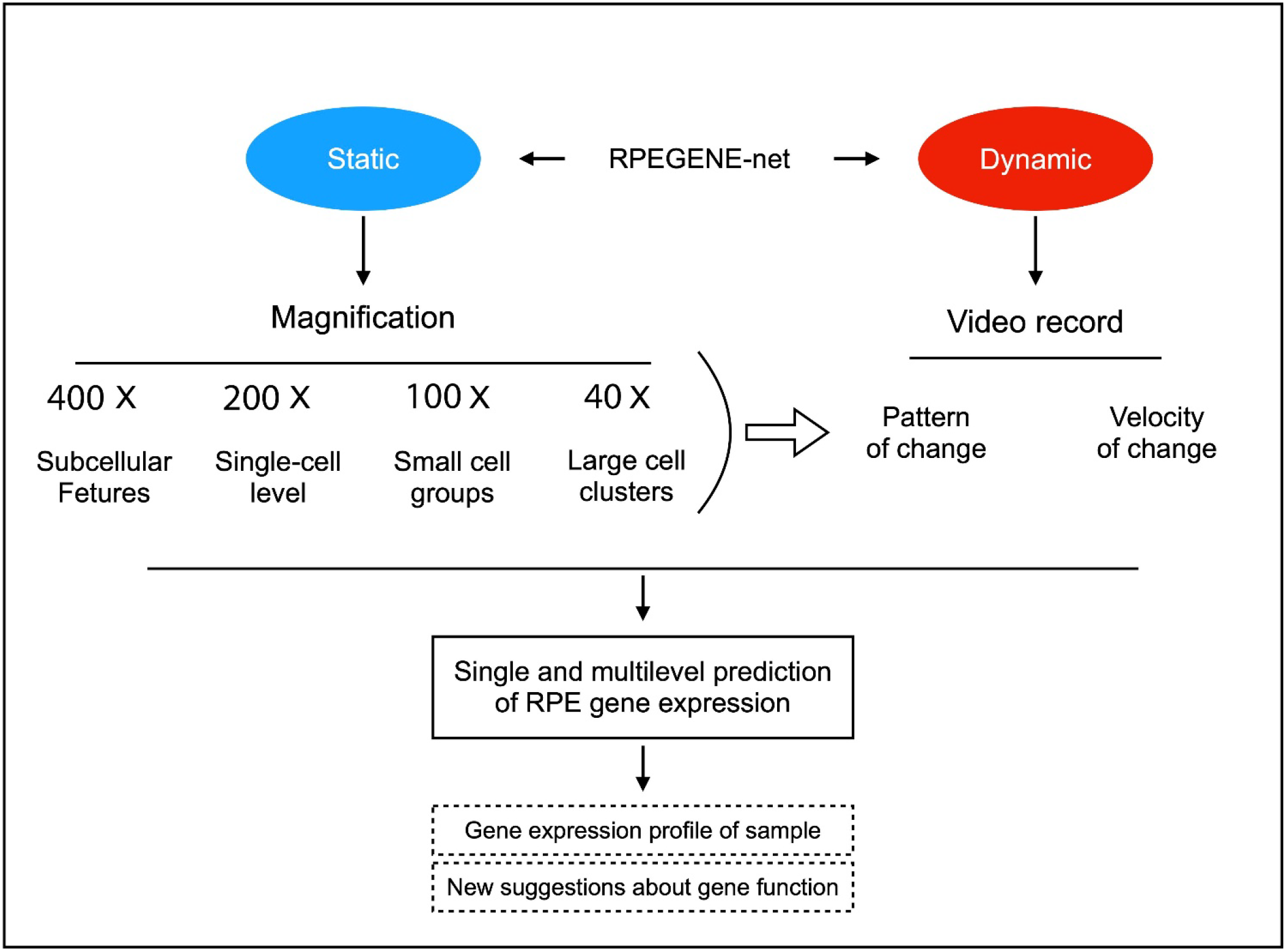
A diagram summarizing the overall research scheme, which involves using different morphological dimensions to determine the best method for gene prediction in a specific cell line. *(In the present article, we only developed the static limb of the software.)*

In the current study, we analyzed twelve models, ranging from convolutional neural networks (CNNs) to advanced transformer-based architectures, to predict the expression of six genes involved in the EMT process of RPE cells (α-SMA, ZEB1, TGF-β, CD90, β-catenin, Snail) and classify five treatment groups (aflibercept, bevacizumab, dexamethasone, aflibercept + dexamethasone, and untreated control). To boost model performance, we trained the algorithms on images captured at different magnifications and integrated their extracted features for final predictions.

### 2.1 Dataset description

For our analysis, we developed a model to predict the expression of six genes—α-SMA, ZEB1, TGF-β, CD90, β-catenin, and Snail—using whole-slide live-cell phase-contrast images of RPE cells. The dataset included a sample derived from an organ donor candidate, which was cultured and expanded to passage 5 before being treated with various drug regimens (detailed below). Microscopic imaging was performed at magnifications of 40x, 100x, 200x, and 400x, with each image captured at a resolution of approximately 1944 × 2592 pixels in TIF format. The experimental procedures are described in detail in the sections below.

#### 2.1.1. RPE cell culture

Human cadaveric eyeballs from a 25-year-old male organ donor were processed for RPE cell culture in a sterile laboratory environment within 48 hours postmortem. Informed consent was obtained from the next of kin for the use of human cadaver tissue in this study. Adnexal tissues were removed, and a limbal incision was made to drain the vitreous. The pigmented inner layer was meticulously dissected into small fragments to initiate RPE cell explant culture. The culture medium consisted of Dulbecco’s Modified Eagle Medium/Nutrient Mixture F-12 (DMEM/F12, Bio Idea, Iran) supplemented with 10% fetal bovine serum (FBS, Gibco, Germany) and 1% penicillin/streptomycin (Shellmax, Iran). To verify the identity of RPE cells, RPE65 expression—a key marker for RPE cells—was assessed via PCR. RNA (100 ng) from both early (1^st^) and late (6^th^) cell passages was analyzed, and RPE65 expression levels were visualized using gel electrophoresis [21]. For subsequent experiments, RPE cells from passages 5 to 7 were used.

The experimental procedure involved dividing RPE cells (106 cells per 10 cm^2^ culture plate) into five groups, each receiving a specific treatment. Group 1 was treated with bevacizumab (312.5 μg/mL, Stivant®, CinnaGen Co., Iran), Group 2 with aflibercept (500 μg/mL, Tyalia®, CinaGene Co., Iran), and Group 3 with dexamethasone (50 μg/mL, Darou Pakhsh Co., Iran). Group 4 received a combination of aflibercept and dexamethasone at specified concentrations, while Group 5 served as the control group with no drug treatment. The drug concentrations were derived from routine clinical dosages for vitreoretinal diseases (2 mg/0.05 mL aflibercept, 1.25 mg/0.05 mL bevacizumab, and 200 μg/0.05 mL dexamethasone, diluted in a 4 mL vitreous cavity volume). Following a 24-hour treatment period, images of the cells were captured for subsequent image analysis, and RPE cells were harvested for gene expression analysis.

#### 2.1.2 Gene expression analysis

RNA extraction from both treated and control groups, cDNA synthesis, and real-time PCR were conducted to evaluate gene expression qualitatively. Total RNA was extracted following the manufacturer’s instructions using the RNeasy kit (Parstous, Iran) and genomic DNA was removed via silica column purification. Single-strand cDNA was synthesized using the Easy cDNA Synthesis Kit RNA (Parstous, Iran). Primer sequences for target genes—α-SMA, ZEB1, TGF-β, CD90, β-catenin, Snail—and the reference gene β-actin were designed using AllelID software (Premier Biosoft International, Palo Alto, CA, USA, v.7.5) (Table 1).

**Table 1.**
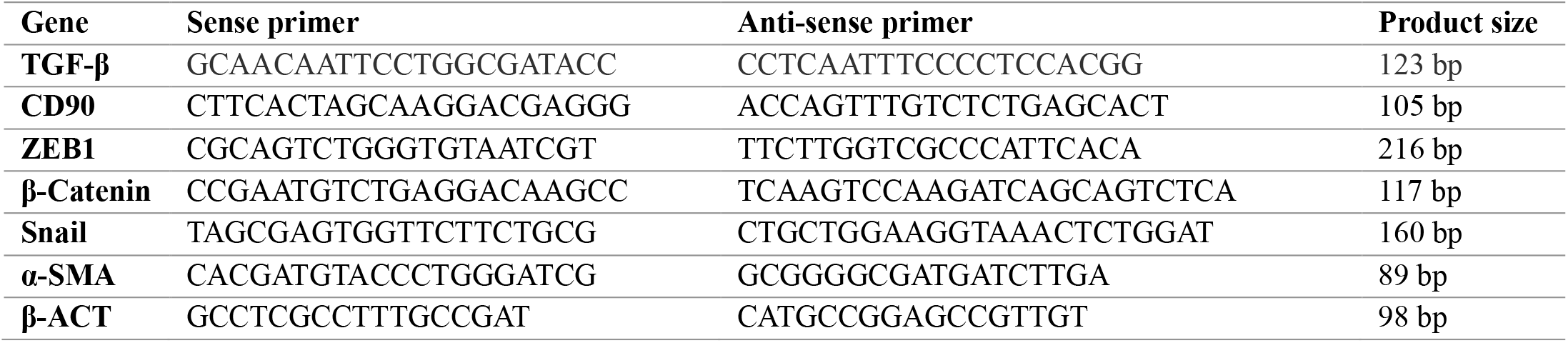
The primer sequences of the genes under study.

Real-time PCR was performed using RealQ Plus Master Mix Green (Ampliqon, Denmark) in a Magnetic Induction Cycler (Mic qPCR, Bio Molecular Systems, Australia). Each PCR reaction consisted of 1.5 μL of forward and reverse primers, 7.5 μL of master mix, and 6 μL of DNA template. The reaction ran for 40 cycles using a one-step thermal profile: a 15-minute hold at 95 °C, followed by a 10-second denaturation phase at 95 °C, and a 45-second annealing and extension phase at 61 °C. The 2^−ΔΔCt^ method was used for the relative quantification of RNA expression.

#### 2.1.3 Image capture

Before mRNA extraction, RPE cells from the treated and control groups were imaged using a JENUS inverted phase-contrast microscope. High-resolution images were captured with a 12-megapixel Microbin camera (Microteb Co., Iran) in TIF format. Images were taken at magnifications of 40x, 100x, 200x, and 400x to facilitate subsequent morphometric analysis. These images constituted the primary dataset for subsequent image processing and the development of analysis algorithms.

### 2.2 Preprocessing of the dataset

Pre-processing is a critical step in quantitative image analysis, addressing limitations inherent to imaging techniques, such as variations in illumination, noise, and artifacts introduced during the image acquisition process. Microscopy images (Figure 2) are particularly prone to these challenges. To mitigate these inconsistencies, we applied vector-based color normalization [22], which effectively normalized the variance in the target images and enhanced their overall quality. As shown in Figure 2, the inconsistency in image intensity was fully resolved following the pre-processing steps.

**Figure 2.**
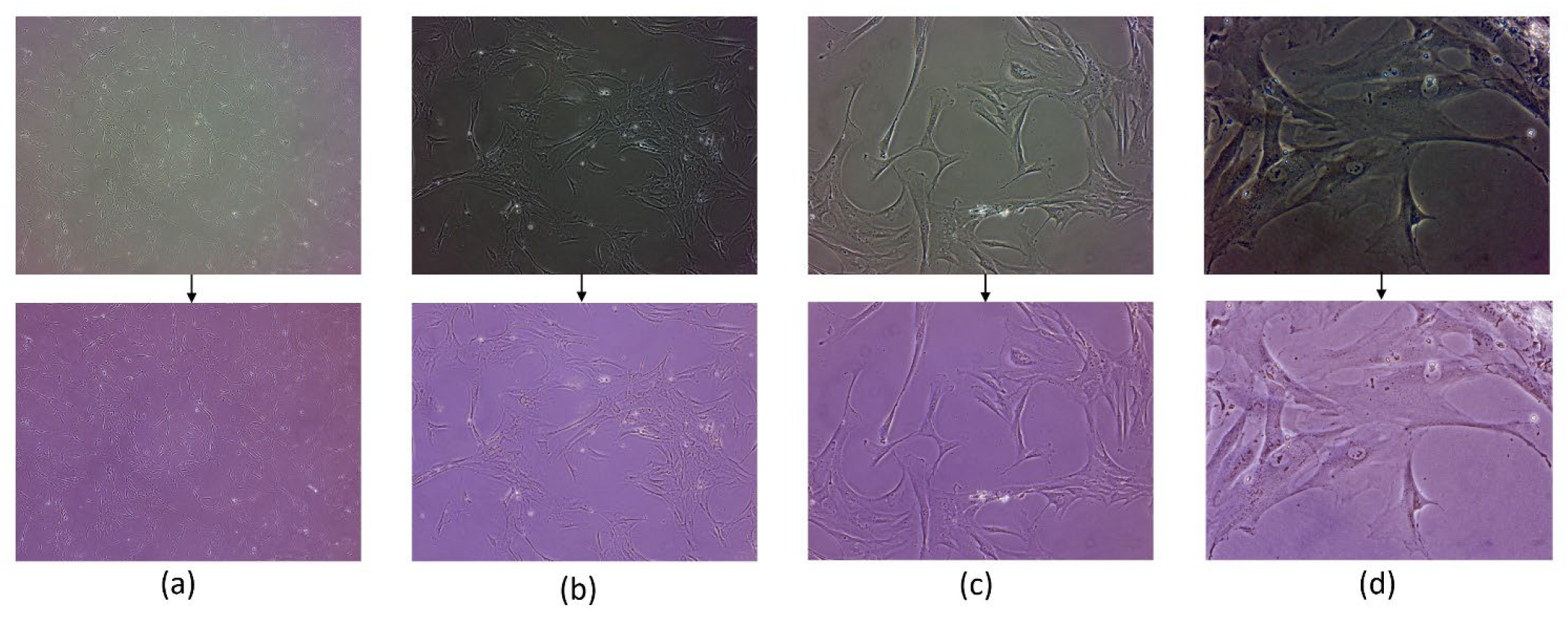
Original sample slide and pre-processed images with magnifications: a) 40x, b) 100x, c) 200x, d) 400x.

#### 2.2.1 Generating image patches

Due to the large size of the original images, we extracted *Np* overlapping patches from the dataset using a fixed patch size and step size. These patches were then organized into a matrix of size *Np × (3 × m × n)*, where *Np* is the number of extracted patches, 3 represents the color channels, and m and n denote the width and height of each patch, respectively. For our experiments, we set *m = n =224 pixels*, yielding approximately 288 overlapping patches per sample (Figure 3). To maximize the utility of information from multiple magnifications, patches were extracted at different scales for each patient. To ensure data quality, we excluded patches containing less than 50% informative pixels, as these could introduce noise or background artifacts.

**Figure 3.**
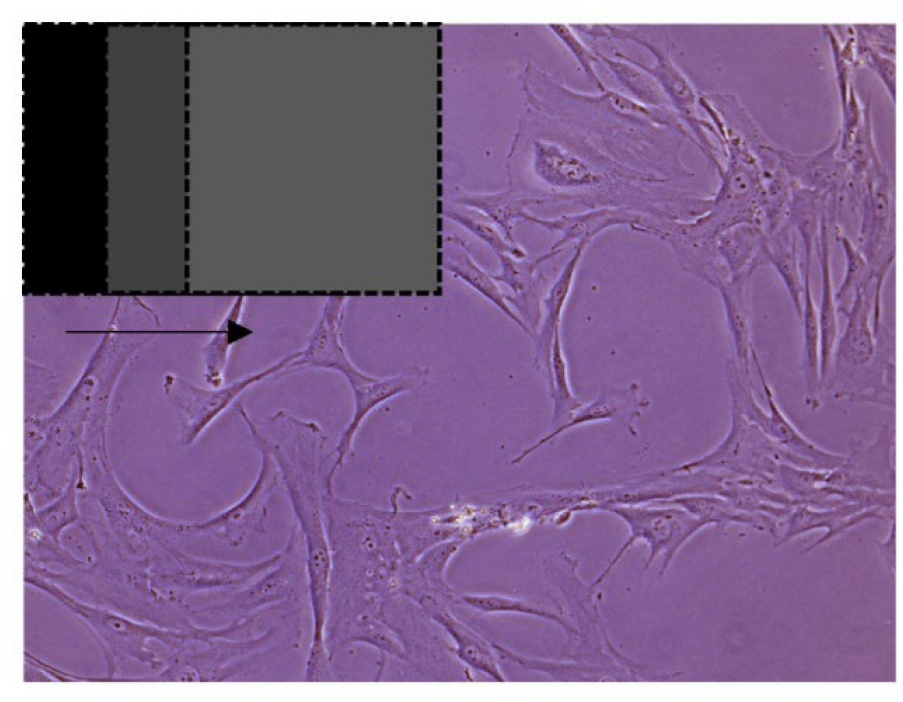
Extracting overlapping patches.

#### 2.2.2 Training, validation and test data splitting

To evaluate our models’ capacity, robustness, and generalization to new images, 20% of the subjects were randomly selected to create an independent test dataset. For the remaining data, we implemented a random training-validation split using 10-fold cross-validation. This method is widely recommended for random splitting as it maximizes dataset utilization, ensuring every data point is used for both training and validation. Such an approach is particularly beneficial when working with smaller datasets.

To prevent overestimation of the model’s performance, it is crucial to avoid using the same image patches from a subject in both training and validation datasets. To address this, we adopted a subject-wise splitting approach, effectively eliminating data leakage between the training, validation, and test datasets. In total, we obtained 12,000 images per magnification for training and 4,100 image patches for testing.

#### 2.2.3 Gene expression values pre-processing

In our study, we investigated six genes associated with EMT process. Alongside image data, the gene expression data also required preprocessing [23, 24]. To prevent model bias towards genes with higher expression levels, we applied a logarithmic transformation to the normalized gene counts (x). To avoid the issue of infinite values resulting from the logarithmic transformation, we added 1 to the gene counts, as shown in Eq. 1. This transformation effectively addressed potential skewness and extreme values in the data: (log_2_(1 + *x*))

#### 2.2.4 Data augmentation

Data augmentation is a widely used preprocessing method in machine learning, particularly when training data is limited. Its primary purpose is to reduce the risk of overfitting by introducing small modifications to the original input images, thereby generating a more diverse dataset. This approach increases both the quantity and variety of training samples. Additionally, data augmentation has the advantage of making the model invariant to specific transformations, enabling it to become less sensitive to minor variations in the input data. This enhances the model’s ability to generalize and perform well on unseen data.

In this study, we employed various augmentation techniques, including geometric and color transformations, on the sample images. For histology image analysis, it is crucial to achieve orientational invariance, similar to how a pathologist examines microscopic images from any angle. To achieve this, we used PyTorch’s torchvision.transforms module to apply random transformations to image patches during training. These transformations included random horizontal and vertical flipping, color jitter, and random equalization, which increased the diversity of the training data. All patches were then converted into PyTorch tensors and normalized to align with the mean and variance of ImageNet-pretrained models.

### 2.3 Gene expression and treatment type prediction models

#### 2.3.1 Construction of the baseline using Autoencoder models

In our research, we addressed two key tasks: predicting gene expression (a regression task) and identifying treatment classes (a classification task). To achieve these objectives, we developed models to classify five distinct treatment classes and predict gene expression across various image patch sizes.

Our analysis incorporated twelve backbone architectures, spanning from CNNs to advanced transformer-based architectures. These included DenseNet versions (DenseNet121, DenseNet169, and DenseNet201) [25], EfficientNet-b5 [26], Inception_v3 [27], RegNet_y_400mf [28], all ResNet versions [29], and Swin Transformer (Swin_b) [30]. For each deep convolutional model pre-trained on ImageNet weights, we implemented a two-stage pipeline to effectively extract relevant features from phase-contrast images.

In the first stage, we fine-tuned each backbone model using genomic spatial image data (magnifications: 40x, 100x, 200x, and 400x) with an autoencoder (AE) mechanism in an unsupervised manner (Figure 4). Autoencoders are particularly effective at extracting essential image features for reliably reconstructing the output image. Leveraging this capability, we ensured the model captured histology-related features crucial for accurate downstream tasks. As shown in Figure 4, the AE model takes an input tensor of size *B* × 224 × *2*24 and outputs a tensor of the same size, where *B* represents the batch size. Then, the pretrained backbones from the AE model were fine-tuned for additional epochs to ensure that the extracted features were histology-relevant and optimized for the classification and regression tasks, which are explained in detail in the following section.

**Figure 4.**
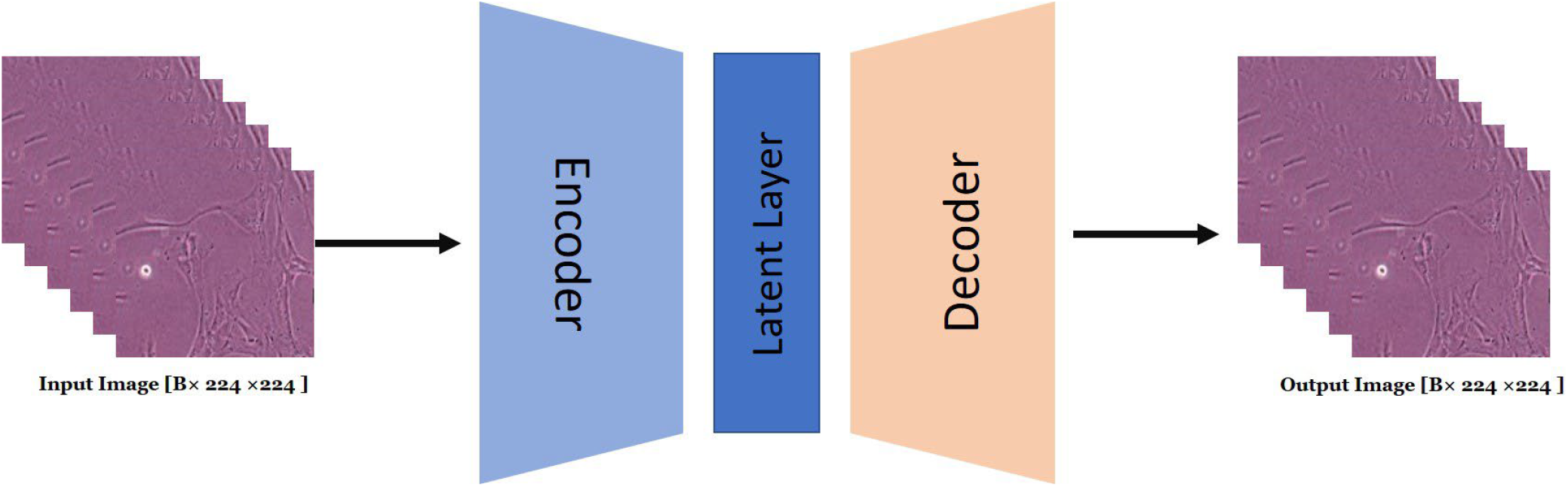
The Autoencoder framework used to create a baseline encoder model.

#### 2.3.2 Gene expression prediction model

Inspired by^24^, the encoders from the first stage were utilized with patches corresponding to every slide, with dimensions of *N*_*p*_×224×224×3, to deliver features with dimensions of *N*_*p*_×1×*N*_*f*_. Here, *N*_*p*_ is the number of patches extracted from a slide, and *N*_*f*_ is the number of outputted features of the model. We applied this procedure to all four corresponding magnifications (40x, 100x, 200x and 400x) related to one slide. To represent the entire slide using a single feature vector, we combined the individual patch-level features to create a vector with dimensions 1 × 1 × *N*_*f*_ as shown in Eq. (1), where *N*_*f*_ represents the number of features after feature aggregation. This process allows us to summarize the slide-level information in a compact and efficient manner as follow:

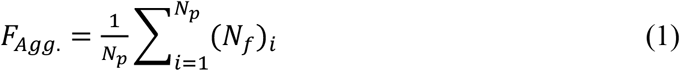

Here, each image patch is represented by a feature vector of size 1 × *N*_*f*_, denoted as (*N*_*f*_)_*i*_, where i is the index of the patch and *F*_*Agg*_ results from features averaging. In other words, the slide-level vector retains the same number of features as the original patch-level vectors and serves as their average representation. This feature is then fed into two 1D convolutional layers with rectified linear unit (ReLU) followed by global average pooling. The first convolution layer uses 256 kernels of size 5×5 and the second one 512 kernels of size 1×1.

To take advantage of the different magnifications corresponding to a single slide image, we developed a feature merging mechanism to extract information about cell morphology from several perspectives, termed multi-stream features, which facilitates a thorough extraction of both inter-cell and intra-cell morphology. To do this, we provide a parallel structure consisting of four networks, each of which extract its own features. We concatenated these outputted features and finally fed them to the output layer with 6 neurons with linear activation functions to jointly train and predict the gene expression related to the slide. We used mean squared error (MSE) as the loss function and optimized the model using Adam with a learning rate of 0.0001 and a decay of 0.01 for each epoch. We set the number of batches and epoch to 16 and 2000 respectively. Despite the limited size of our dataset, we implemented several strategies to prevent overfitting and ensure model robustness. A 10-fold cross-validation strategy was employed to ensure model performance was not overly reliant on a single data subset. Overfitting was further mitigated through various techniques, such as data augmentation during the training process.

Additionally, to minimize model parameters, we used 1D convolutional layer to have an appropriate model parameter compared to the data. Block Diagram of proposed prediction model are shown in Figure 5.

**Figure 5.**
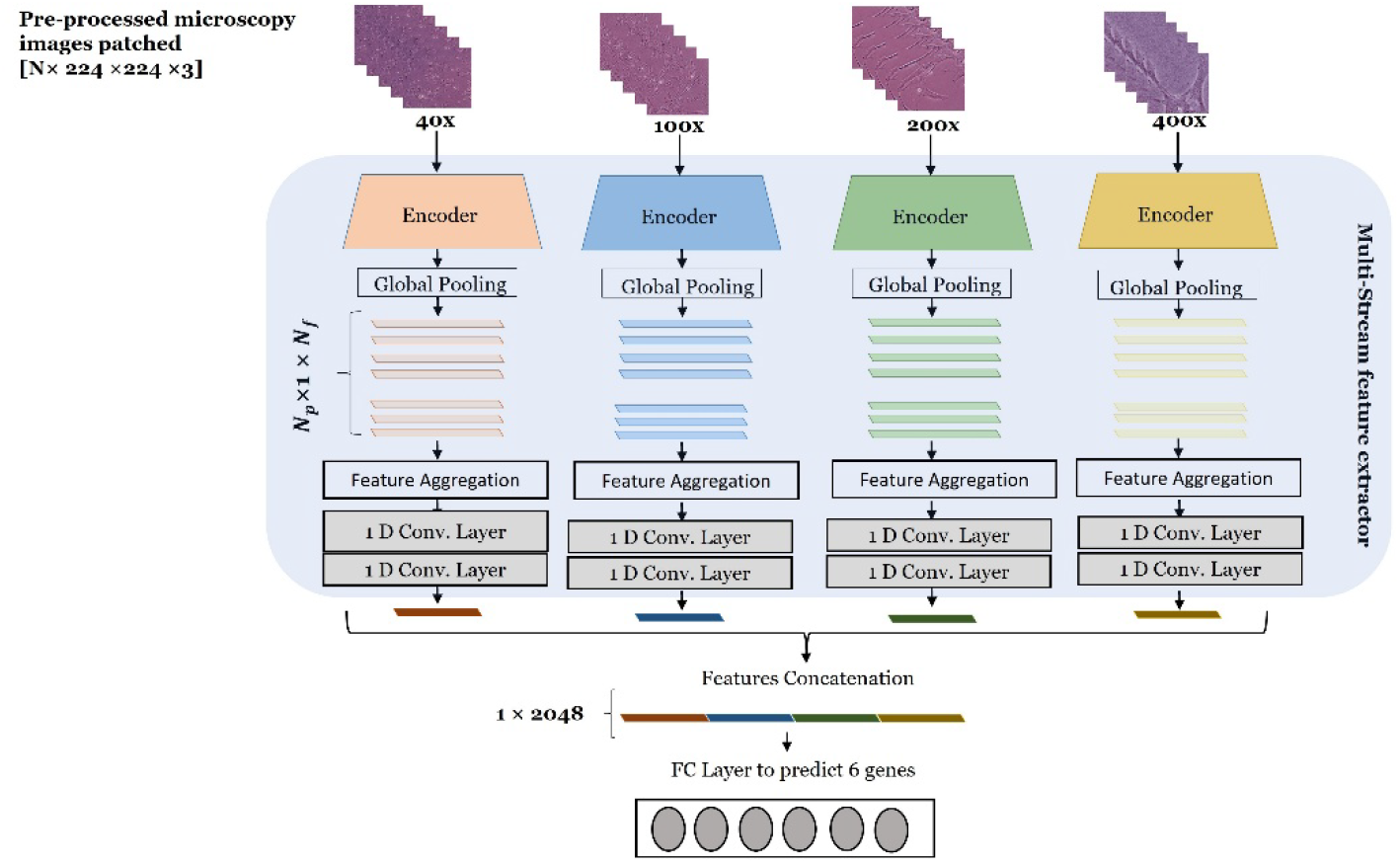
Block diagram of our proposed method to predict gene expressions.

As demonstrated in Figure 5, during the testing stage, our model aggregates the predicted gene expression of each magnification to obtain the final slide-level gene expression prediction.

#### 2.3.3 Treatment type prediction model

The baseline model architecture is modified in the way that be appropriate for a classification task to distinguish treatment classes. As illustrated in Figure 4, encoders from the AEs are utilized to extract relevant features from each magnification level. After feature of each magnification extracted, we concatenate those multi-stream features to feed them to the fully connected layer to output 5 classes of treatment. To optimize the classification model, we employed the Adam optimizer, Cross-entropy loss function, and a learning rate of 0.0001 with a decay of 0.01 per epoch. These settings were used to update the model’s weights during training. A batch size of 16 samples was used for each training step. Furthermore, a sigmoid activation function is applied to the final linear layer to produce probability scores for predicting one of five treatment classes. Figure 6 visually demonstrates our classification approach.

**Figure 6.**
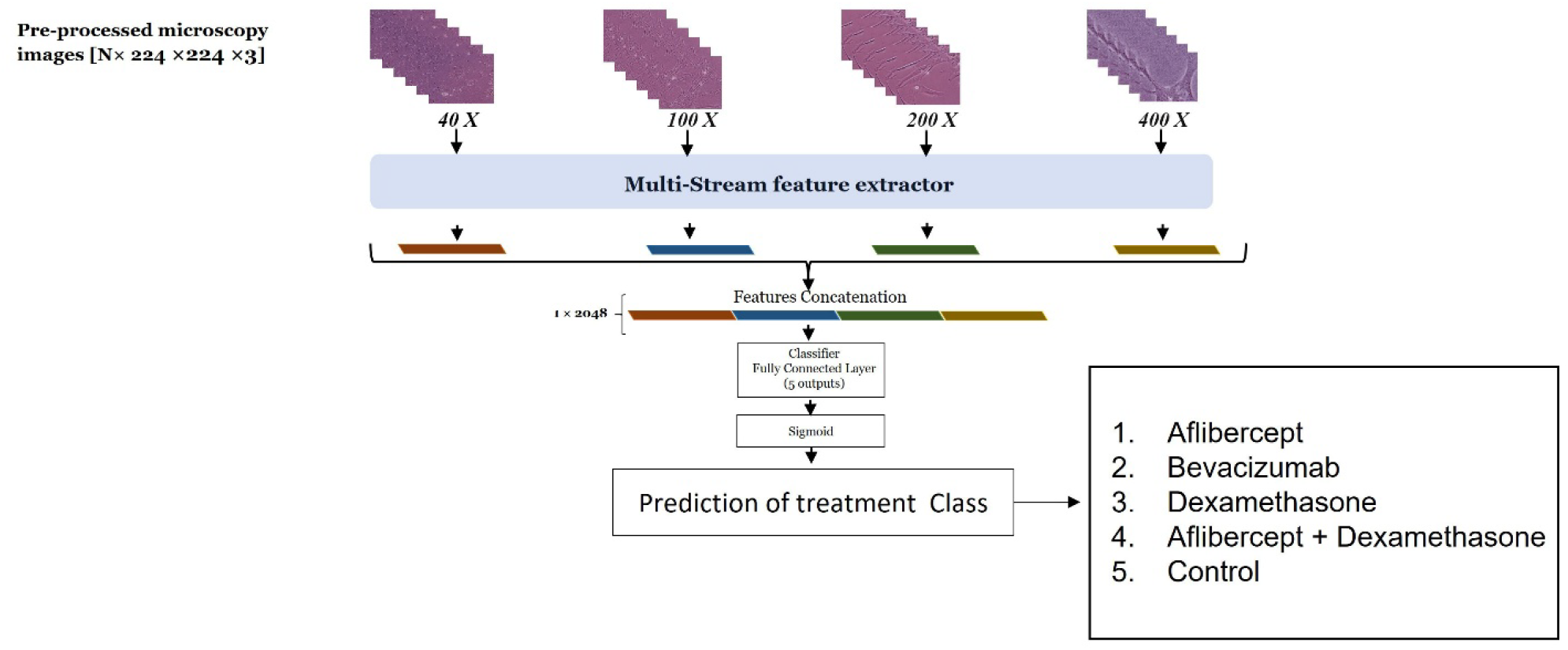
Block diagram of the classification model used to predict the treatment class.

## 3. Evaluation of the Models

### 3.1 Evaluation metrics for regression model

As mentioned earlier, predicting gene expression levels is a regression task, while identifying treatment class is a classification task. The performance of the regression model was evaluated using MAE, RMSE, and the Pearson Correlation Coefficient (PCC):

MAE and RMSE measure the difference between predicted and actual gene expression values. The R2 score indicates how well the model fits the data, with higher scores indicating a better fit:

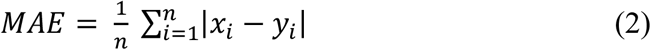

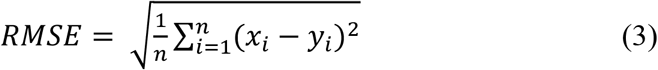

where n is the total number of samples, *x*_*i*_ is the target value of sample i, and *y*_*i*_ is the predicted value. Both error measures yield an average prediction error of a model, ranging from 0 to ∞, with smaller values indicating more accurate predictions.

The PCC was used to determine how well gene expression predictions from sample images matched actual gene expression levels. It measures the linear relationship between two variables, with scores ranging from −1 to 1. A score of 1 indicates a perfect positive relationship, −1 indicates a perfect negative relationship, and 0 indicates no relationship.

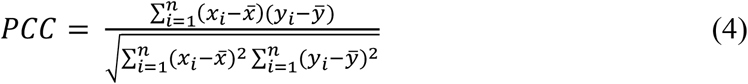

where *x*_*i*_ and *y*_*i*_ are the target and predicted genes, respectively, 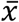 is the mean of the *x*_*i*_, and 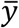is the mean of the *y*_*i*_.

### 3.2 Evaluation Metrics for Classification Model

The following metrics were used to evaluate the treatment classification model: Sensitivity, Specificity, precision, Accuracy, and F-score:

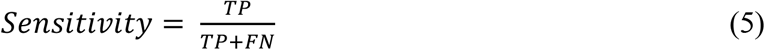

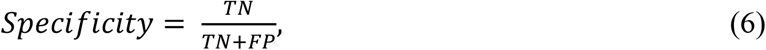

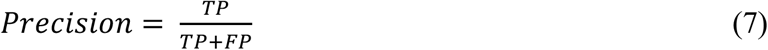

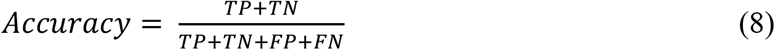

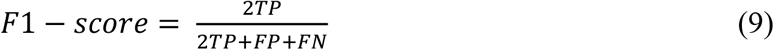

## 4. Results

Our models were implemented using PyTorch and Python 3.9, and training was performed on an RTX 4060 Ti GPU. We trained and evaluated a total of 12 models on our dataset for two tasks: distinguishing treatment classes into five groups (referred to as groups 1 to 5, as described in the “RPE Cell Culture” section) and predicting the expression of six genes.

The models were based on pre-trained networks, including DenseNet121, DenseNet169, DenseNet201, EfficientNet-b5, Inception_v3, RegNet_y_400mf, all ResNet versions, and Swin_b. All reported results were obtained by evaluating the models on an independent test dataset, ensuring unbiased performance metrics.

### 4.1 The results of regression models to predict the gene expression

The median PCC values for each gene from the held-out test set, using 12 different deep learning backbones, are presented in Table 2. The values highlighted in bold represent the optimal performance (i.e., the highest PCC) for each gene. Notably, DenseNet121 demonstrated the best performance in 4 out of 6 gene expressions (α-SMA, ZEB1, TGF-β and Snail), demonstrating high predictive accuracy. Meanwhile, ResNet34 excelled in predicting the remaining two genes (CD90 and β-Catenin), exhibiting strong correlations between predicted and target values.

**Table 2.**
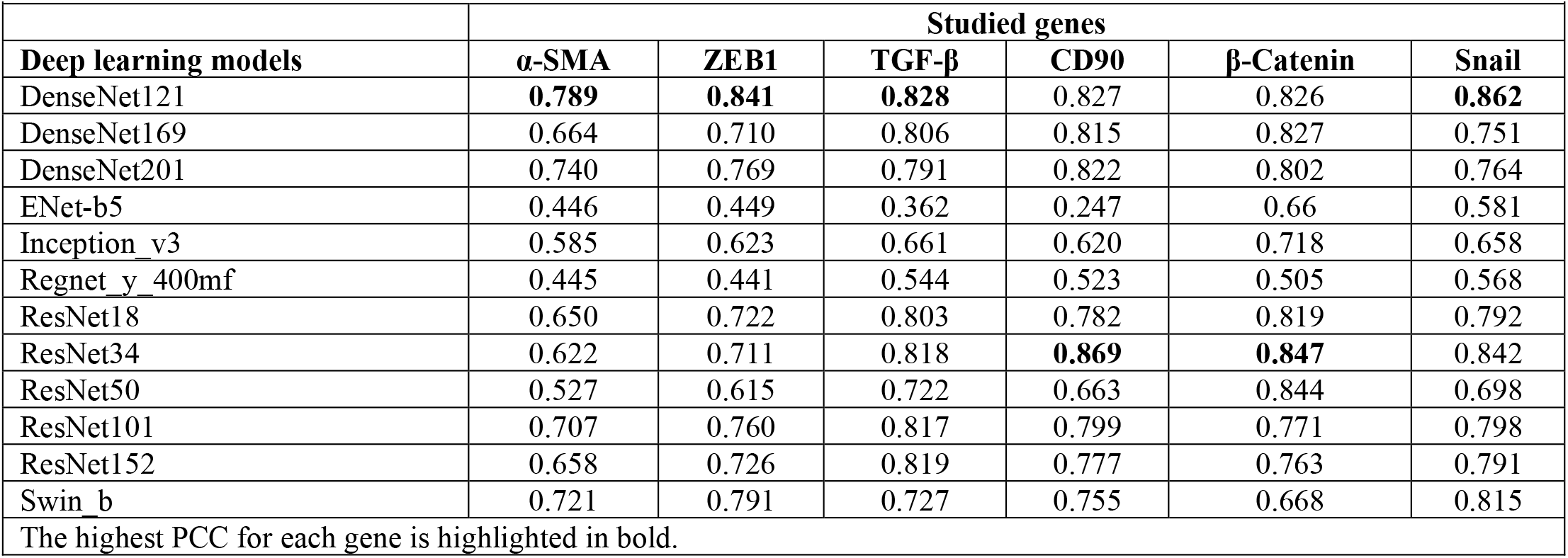
The median Pearson Correlation Coefficient (PCC) of 6 predicted gene expressions using 12 pre-trained models.

To accurately estimate the predicted mean and standard deviation of gene expressions, we randomly selected three slides from each treatment class. Multiple patches were then extracted from each slide, and the gene expression values for each patch were predicted using the DenseNet121 model. The average predicted gene expression values for the patches were calculated to assign a single value to each slide. Finally, the mean and standard deviation of these slide-level values were calculated to represent the final prediction of gene expression for each treatment class. Figure 7 compares the predicted and actual gene expression levels across the five treatment groups.

**Figure 7.**
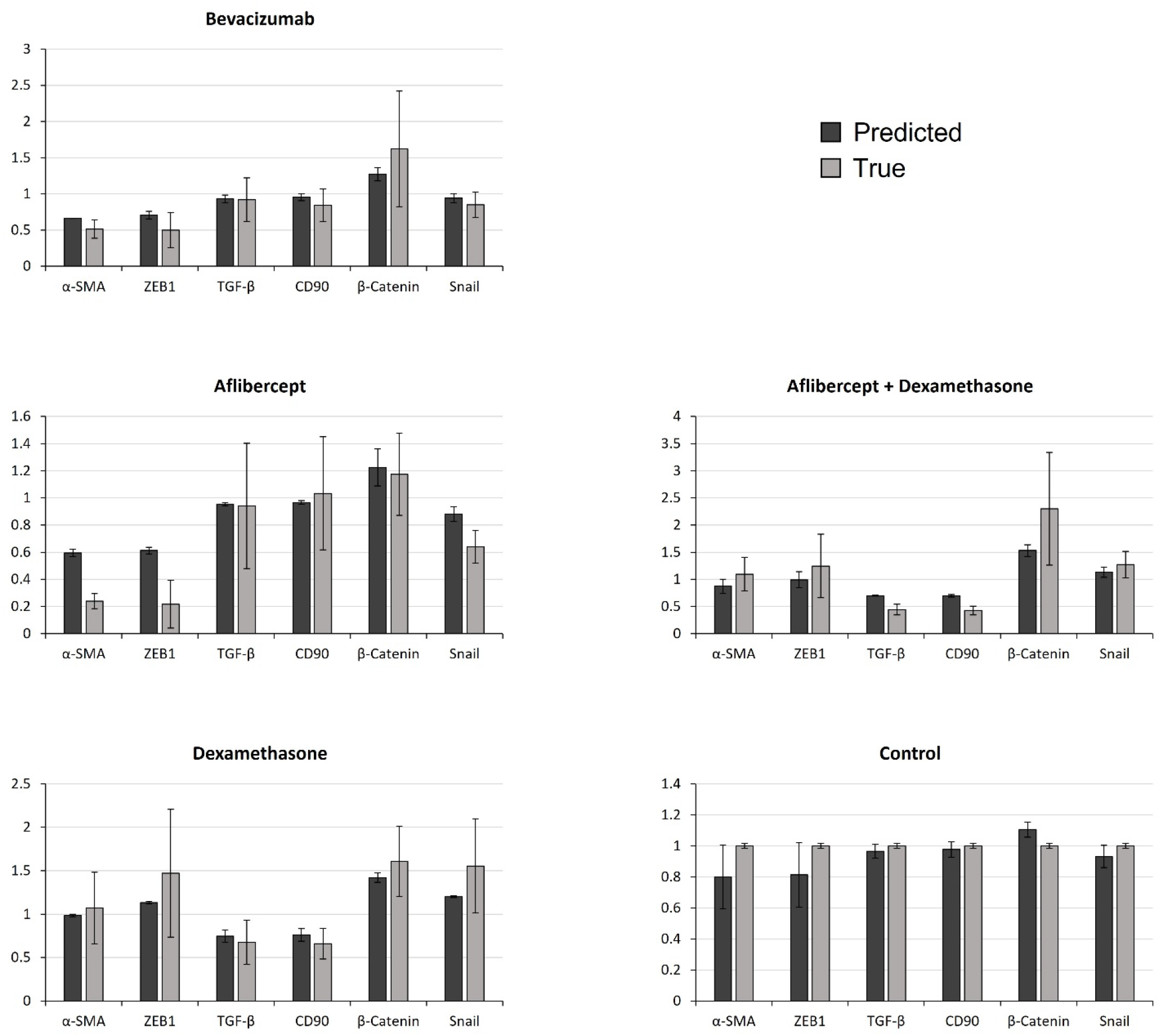
Comparison between the predicted and true levels of gene expression across the five treatment groups.

Additionally, we evaluated the performance of our proposed method by visualizing the distribution of gene expression values using a violin plot, as shown in Figure 8. This visualization highlights the spread and density of gene expression predictions, providing further insights into the model’s performance.

**Figure 8.**
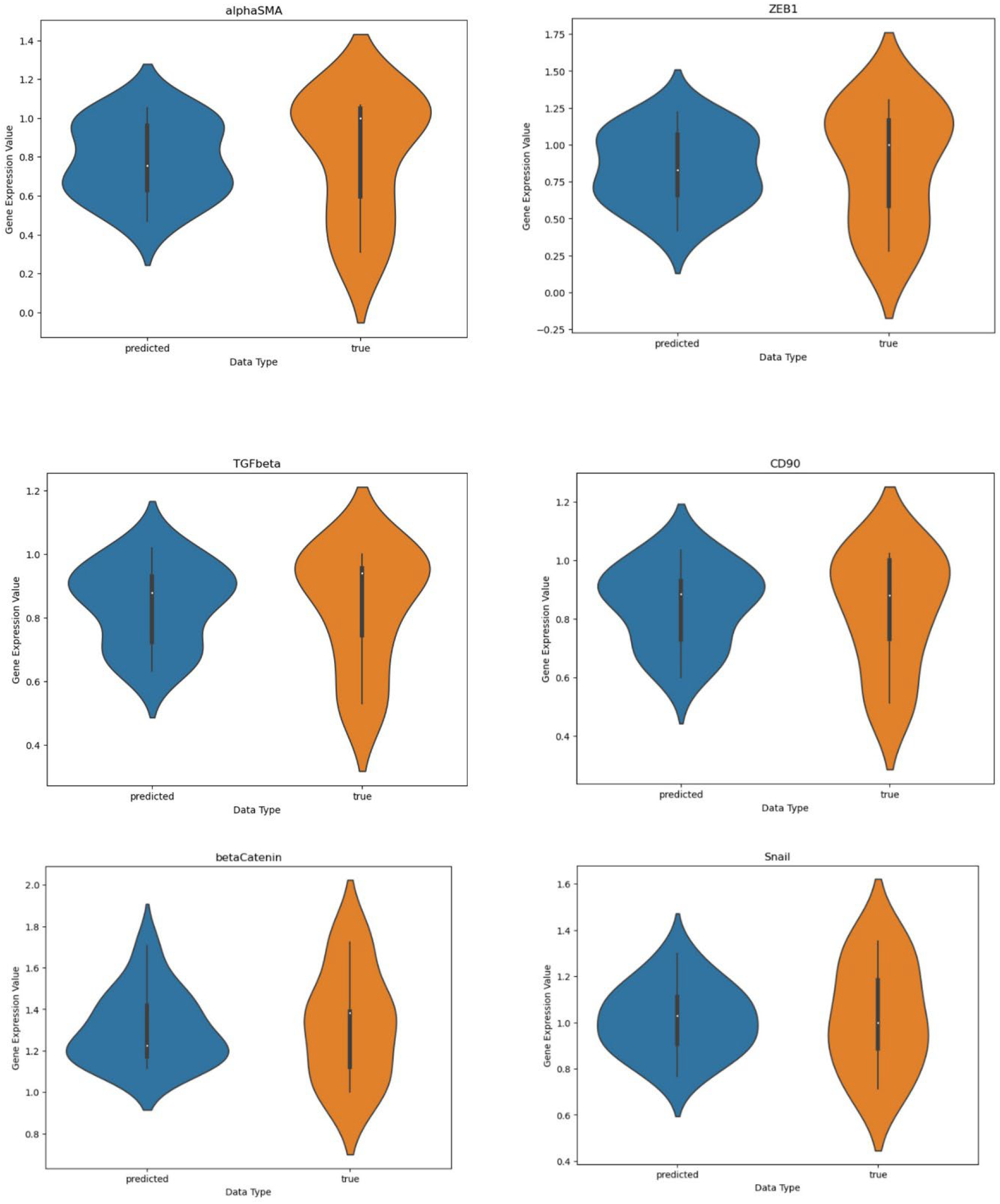
Sample true vs. predicted gene expression violin plots for DenseNet121.

For a more detailed analysis, we assessed the performance across four magnifications (40x, 100x, 200x, and 400x). The results of the DenseNet121 model for each magnification and the combined multi-magnification approach are summarized in Table 3. To compare performance across magnifications, we calculated the R^2^ score. As shown in Table 3, using all four magnifications together resulted in better performance, demonstrating the effectiveness of the multi-magnification approach.

**Table 3.** Comparison of R^2^ Scores for Different Genomic Spatial RPE Input Patch Magnifications.

|  |  |  |  |  |  |  |
| --- | --- | --- | --- | --- | --- | --- |
| Genes<br>Different spatial<br>Genomics patch | alphaSMA | ZEB1 | TGFbeta | CD90a | betaCatenin | Snail |
| 40 x | **0.5239** | 0.5880 | 0.6248 | 0.6079 | 0.5801 | 0.5311 |
| 100 x | 0.1389 | 0.2170 | 0.2803 | 0.2263 | 0.3584 | 0.1367 |
| 200 x | 0.3061 | 0.3369 | 0.0544 | 0.0726 | -0.1196 | 0.2958 |
| 400 x | 0.3736 | 0.4686 | 0.0496 | -0.0316 | 0.2870 | 0.5106 |
| 4x+10x+20x+40x | 0.4987 | **0.6009** | **0.7413** | **0.6986** | **0.6217** | **0.6719** |

### 4.2 The results of classification model to classify the treatment classes

We utilized the best-performing model from the regression task to classify the treatment classes. Given that DenseNet121 achieved the highest accuracy in predicting gene expression, it was selected for retraining to perform the classification of treatment types. Accordingly, DenseNet121 achieved exceptional performance in the classification of treatments into five classes using an independent test dataset. These results highlight DenseNet121’s effectiveness and reliability in this multi-class classification task as shown in Table 4.

**Table 4.** The results of treatment type classification using DensNet121 Architecture.

|  |  |  |  |  |  |
| --- | --- | --- | --- | --- | --- |
| Criteria<br>Treatment type | Precision (%) | Recall(%) | F1 score(%) | Specificity(%) | Accuracy per class(%) |
| Aflibercep | 100 | 100 | 100 | 100 | 100 |
| Bevacizumab | 100 | 66.67 | 80.00 | 100 | 93.33 |
| Dexamethasone | 100 | 100 | 100 | 100 | 100 |
| Aflibercept +<br>Dexamethasone | 100 | 66.68 | 80.00 | 100 | 93.33 |
| Untreated control | 60.0 | 100 | 75.00 | 83.33 | 86.66 |
| Average | 91.99 | 86.66 | 97.00 | 86.99 | 94.66 |

## Discussion

In this study, we combined phase-contrast microscopy imaging with gene expression measurements obtained via qPCR to develop a deep learning model capable of predicting gene expression levels directly from live-cell images, eliminating the need for staining or fixation. By categorizing cells into distinct groups and exposing them to specific drugs, we aimed to induce the expression of genes associated with cell morphology. We captured images at multiple magnifications (40x, 100x, 200x, and 400x) and successfully trained the system to estimate the expression of key genes involved in the process of EMT— α-SMA, ZEB1, TGF-β, CD90, β-catenin, and Snail—with high precision and sensitivity. These genes are critical in RPE pathology, particularly in retinal diseases such as proliferative vitreoretinopathy (PVR), where EMT drives fibrosis and vision loss [18, 19].

Our study introduces RPEGENE-Net, a novel deep learning framework that predicts gene expression patterns directly from microscopy images of RPE cells. This approach bypasses traditional gene expression analysis techniques, offering a non-invasive, cost-effective alternative. RPEGENE-Net demonstrated robust performance, as evidenced by high Pearson correlation coefficients and low MAE metrics, underscoring its potential for both research and clinical applications.

To ensure the reliability of RPEGENE-Net, we conducted an extensive evaluation of diverse model architectures for gene expression prediction. Our analysis encompassed twelve models, spanning from CNNs to advanced transformer-based architectures. To further enhance the model’s performance, we employed transfer learning with a novel two-step adaptation process. Initially, we trained an autoencoder on our dataset to capture a compact, data-specific representation. The encoder from this AE was then fine-tuned for the gene expression prediction task. This innovative approach ensured that the model’s weights were tailored to the unique characteristics of our dataset, resulting in highly accurate and reliable predictions.

Among the evaluated architectures, DenseNet and ResNet emerged as the most effective for our prediction task. As shown in Table 2, DenseNet121 excelled in predicting genes such as α-SMA, ZEB1, Snail, and TGF-β, while ResNet34 outperformed in predicting CD90 and β-Catenin. DenseNet’s densely connected layers promote efficient feature reuse, enabling the model to capture comprehensive and distinctive patterns in the data. This characteristic, combined with its relatively lightweight structure, contributed to its superior performance compared to other architectures. The dense connections facilitate robust extraction of high-level features, which are critical for accurate gene expression prediction.

Similarly, ResNet’s residual connections effectively address the vanishing gradient problem, allowing deeper architectures to maintain strong performance without degradation. However, both DenseNet and ResNet architectures showed diminished performance when using deeper variants, likely due to the limitations of the dataset size. In general, deeper networks tend to benefit from much larger datasets, highlighting the need for more extensive data in future studies to fully harness the potential of these architectures.

Artificial intelligence (AI) tools are invaluable for analyzing cell images and predicting gene expression [12]. Unlike previous studies that relied on stained cell images or processed slides [31], our approach predicts gene expression patterns using live-cell images captured under phase-contrast light microscopy— without the need for any experimental staining or cell processing.

The rationale for using different magnifications in cell imaging was to provide a comprehensive dataset enriched with bioinformatics insights across various biological scales. At 400x magnification, we focused on subcellular structures; at 200x, we captured single-cell details; at 100x, we examined interactions between a few cells; and at 40x, we assessed characteristics of cell populations. As shown in Table 3, the highest coefficient of determination (R^2^) was achieved by combining data from four different magnifications, compared to using any single magnification alone. This finding supports our hypothesis that a multilevel approach to cellular imaging provides the most accurate prediction of gene expression for a given sample. This observation may stem from the multifunctionality of each gene, which is reflected in its role across various cellular and subcellular structures, as well as in cell-to-cell interactions. In our study, all genes analyzed showed strong relevance to cell population images (40x magnification), while α-SMA, ZEB-1, and Snail were also prominently associated with subcellular structures (400x magnification) (see Table 3). This finding may reflect the role of each gene in causing morphological changes at the cellular or subcellular level.

α-SMA is a well-known marker of myofibroblasts, which are crucial in wound healing and tissue remodeling [32]. In the context of the EMT of RPE cells, α-SMA indicates a more contractile phenotype, significant in the development of fibrotic diseases like PVR [33]. These morphological changes, including stress fiber formation and increased cell contractility [34], are more discernible at lower magnifications (40x), where overall cell shape and interactions are visible. Conversely, genes like ZEB1, TGF-β, CD90, β-catenin, and Snail, involved in EMT, have expression patterns more dependent on subcellular structures and finer morphological details, better observed at higher magnifications (400x). ZEB1 and β-catenin are transcription factors involved in nuclear processes [35, 36], while TGF-β and CD90 influence fine cytoskeletal remodeling [37, 38]. This distinction highlights the importance of the cellular context and the observation scale in understanding gene function and expression patterns. Thus, α-SMA is better captured at lower magnifications, while subcellular details of other EMT-related genes are clearer at higher magnifications, emphasizing the need for multilevel imaging approaches.

In addition to the insights gained from the known functions of the gene in question, image analysis at different magnifications can provide further clues about the gene’s involvement in various aspects of cellular behavior. For instance, in our case, a higher R^2^ value observed at 40x magnification emphasizes the gene’s potential role in cell-cell interactions, as reflected in the organization and patterns of cell populations. Conversely, a greater R^2^ value at 400x magnification underscores the gene’s significant contribution to intracellular responses.

The findings of this study offer several clinical implications. First, the RPEGENE-Net model can be utilized to predict gene expression in RPE cells in vitro. While not as precise as PCR analysis, it provides a cost-effective alternative for gene analysis in experimental studies. However, to enhance the model’s accuracy and utility, it will require training with larger and more diverse gene expression datasets. Second, RPEGENE-Net could serve as a quality control tool during specific stages of RPE cell preparation for transplantation, particularly in situations where RPE preservation, cost efficiency, and time constraints are critical considerations. Beyond its application to RPE cells, this study introduces a simple yet effective approach for predicting gene expression from cell images. This method holds broader potential, such as its application to other cell types (e.g., blood cells for cancer detection) or even in vivo cell imaging as a rapid screening tool.

### Powers and limitations

The present study offers two key innovations. First, it introduces a deep learning approach for predicting gene expression in *RPE cells in vitro*. Second, it employs an AI-based strategy combining multi-level magnification and deep learning integration of multi-scale data. While the results are promising, the relatively small dataset size poses a limitation. To address this challenge, we implemented several strategies. One involved extracting image patches to significantly expand the dataset. Data augmentation not only increased the dataset size but also enhanced the model’s generalization and robustness. Future research could focus on further expanding the dataset to improve model performance. Moreover, extending the applicability of RPEGENE-Net to other cell types and disease models would be a valuable direction for exploration.

## Conclusions

This proof-of-concept study demonstrated that integrating RPE cell images captured at different magnifications—ranging from subcellular structures to large cell clusters—significantly enhances the predictive capability of models for determining the gene expression profiles of specimens. This approach may have clinical implications for the quality control of cell-based products and could also facilitate the discovery of novel gene functions through morphological analysis. Among the models tested in this study, the DenseNet121model exhibited the best performance for this purpose.

## Ethics Approval and Consent to Participate

The protocol used in this study was approved by the Ethics Committee of Shiraz University of Medical Sciences (IR.SUMS.REC.1401.458). Informed consent was obtained from the next of kin for the use of human cadaver tissue in this study.

## Consent for publication

Not applicable

## Competing interests

The authors report no commercial or proprietary interest in any product or concept discussed in this article.

## Funding

This study was supported by Shiraz University of Medical Science (Grant # 26160).

## Author Contributions

MH.N. and F.SJ. were involved in the project design, obtaining the data, and performing the experiments. T.M. and N.T. were involved in machine learning and deep learning analysis. All authors were involved in writing the manuscript and revising the final version.

## Acknowledgments

The authors would like to thank the directors of Shiraz University of Medical Sciences for supporting this research.

## Data availability

The data used to support the findings of this study are available from the corresponding author upon request.

## Notes

### Competing Interest Statement

The authors have declared no competing interest.

